# *Ex vivo* primary liver sections recapitulate disease phenotype and therapeutic rescue for liver monogenic diseases

**DOI:** 10.1101/2023.03.23.533840

**Authors:** Dany Perocheau, Sonam Gurung, Loukia Touramanidou, Claire Duff, Garima Sharma, Neil Sebire, Patrick F. Finn, Alex Cavedon, Summar Siddiqui, Lisa Rice, Paolo G.V. Martini, Andrea Frassetto, Julien Baruteau

**Author notes:** **Corresponding author:** Dr Julien Baruteau, Genetics & Genomic Medicine Department, Great Ormond Street Institute of Child Health, University College London, 30 Guildford Street, WC1N 1EH. London.

## Abstract

In academic research and the pharmaceutical industry, *in vitro* single cell line cultures and *in vivo* animal models are considered as gold standards in modelling diseases and assessing therapeutic efficacy. However, both models have limitations, with incomplete reproduction of pathophysiological characteristics and absence of 3-dimensional architecture with cell lines or the use of live animals brings ethical considerations, limiting the experimental scale and design. The use of precision-cut tissue slices can bridge the gap between these mainstream models as this technique combines the advantages of studying all cell sub-types whilst preserving the tissue-matrix architecture, thereby closely mimicking a mini-organ. Here, we describe an optimised and easy-to-implement protocol for the culture of sections from mouse livers. We show that precision-cut liver sections can be a reliable model for recapitulating the biological phenotype of inherited metabolic diseases, exemplified by common urea cycle defects citrullinemia type 1 and argininosuccinic aciduria, caused by argininosuccinic synthase (ASS1) and argininosuccinic lyase (ASL) deficiencies respectively. Therapeutic response to gene therapy such as messenger RNA replacement delivered via lipid nanoparticles can be monitored, demonstrating that precision-cut liver sections can be used as a preclinical screening tool to assess therapeutic response and toxicity in monogenic liver diseases.

## Introduction

In research and development, isolated primary cells and cell line cultures are models of choice for *in vitro* studies due to their easy access, cost effectiveness and limited effort required for maintenance. However, limitations of modified physiology and loss of differentiation rapidly observed in the absence of tissue specific microenvironment, limit these advantages [1,2]. To mitigate this, *in vitro* three-dimension induced pluripotent stem cells-derived spheres or miniorgan [3] and whole-organ bioreactors from decellularized scaffold, which enable the association of different cell types to recreate a tissue architecture *in vitro* [4], are attractive, especially for therapeutic screening. However, these techniques can be technically challenging and costly to maintain and reproduce, limiting their use for high-throughput screening. Transgenic animals can recapitulate the pathophysiology of monogenic disorders but present their own limitations *i.e*. high cost of maintenance and limitations of experimental design for therapeutic screening due to ethical concerns as outlined by guidelines on animal research [5].

In this context, the precision-cut tissue slice (PCTS) model is a cost-effective and relatively less labour-intensive system hence a valuable compromise, which allows high-throughput testing whilst preserving the tissue architecture [6, 7]. PCTS fills a gap between *in vitro* cell studies and *in vivo* animal research, overcoming major disadvantages of both models. PCTS mimics a mini-organ model, preserving the 3-dimensional aspect of neighbouring cells and extracellular matrix. The development of improved tissue slicers, *e.g*. vibratome [8], has allowed a transition from manually cut slices characterised by heterogeneous width and poor survival rate to reproducible thinner slices with better preserved structural integrity, further facilitated the optimisation of culture conditions such as appropriate oxygen and nutrients. PCTS has been generated from various organs such as liver [9], intestines [10, 11], colon [10], brain [12, 13], lung [7, 14, 15], kidney [16, 17], spleen [18, 19], heart [20, 21] and tumours [22, 23]. They can also originate from various animal models such as mouse [9], rat [24, 25], pig [26] and human surgical wastes [22, 27, 28]. The ease and reproducibility in preparing PCTS facilitate high-throughput therapeutic screening and toxicology studies [10, 16, 29]. Although PCTS require the use of animals implying ethical limitations, the organ from one animal can generate multiple PCTS, thereby reducing drastically the number of animals in agreement with the NC3Rs guidelines [30] and limiting interindividual variations. The PCTS system also avoids the bias of off-target effects.

Chronic liver diseases represent a healthcare burden affecting more than 1.5 billion persons worldwide, causing one of the top five causes of death in working adults [31], with an increase of incidence of 13% over the last 20 years [32]. Some of these diseases require liver transplantation, which can, for some of them, be progressively disregarded in favour of rapidly expanding innovative gene therapy strategies [33]. Developing liver PCTS is an appealing strategy to model a disease phenotype and test preclinical therapeutic effect and potential toxicity in a high-throughput manner.

Here, we present an optimised method for the preparation and culture of precision-cut liver slice (PCLS) with survival of up to 5 days. We show that PCLS recapitulate *in vitro* the biological phenotype of two rare liver inherited metabolic diseases affecting the urea cycle, citrullinemia type 1 caused by argininosuccinate synthase (ASS) deficiency and argininosuccinic aciduria (ASA) caused by arginosuccinic lyase (ASL) deficiency. We then show that PCLS effectively support the proof of concept of gene therapy by rescue of the ASA phenotype by *hASL* mRNA encapsulated in lipid nanoparticles.

## Results

### Key aspects for the preparation and culture of PCLS

Protocols for PCLS preparation and culture vary significantly in the literature with a lack of standardisation, especially for slicing equipment, medium culture, and engineering system. Optimisation for each protocol is key and varies noticeably depending on the tissue of interest. We set up a rapid and optimised protocol to generate PCLS for high-throughput (Figure 1A). At harvest, perfusion of the animal was purposely omitted to ensure a rapid processing of the organ and prevent organ damage as this was proven not to provide any additional benefit for the PCLS model [34]. The liver was extracted in less than 30 seconds following incision and immediately placed in ice-cold organ-protective buffer *e.g*. Krebs buffer [35, 36]. Although slicing fresh liver tissue without embedding has been previously described [9], we embedded the liver in low-melting agarose [37] (Figure 1B) combined with an organ-protective buffer (Figure 1C) to enable optimal cutting conditions on the vibratome, reducing tissue damage and increasing reproducibility in section thickness. Tissue thickness is critical as thin sections allow more cell layers to access nutrients and oxygen [38], reducing cell death. However, sections thinner than 200 μm become difficult to cut homogeneously and can show increased oxidative stress [25]. Slices thicker than 400 μm show low penetration rate of nutrients or therapeutic particles [25, 29]. Therefore, based on PCLS appearance, effects on texture and ease of cutting [38, 39], we opted for a PCLS thickness of 250 μm, producing up to 30 sections per liver lobe of an adult mouse (Figure 1D). The sections were incubated in a liquid-air interface using an insert (Figure 1E) and incubated with 5% CO2 and 21% O2 at 37°C on a shaker. Sections had to be incubated in culture medium ideally within 90 minutes to up to 3 hours following harvest after which there was rapid occurrence of cell death [29]. Supplemented media with glucose, serum and insulin has been described with potential benefit in preserving the viability and functionality of slices [40]. Insulin could facilitate PCLS culture by improving hepatocyte metabolism [41, 42] but we kept purposely a higher glucose concentration in our culture medium to maintain viability, as described by others [25, 42, 43]. The optimised culture medium used was replaced every 2 days.

**FIGURE 1.**
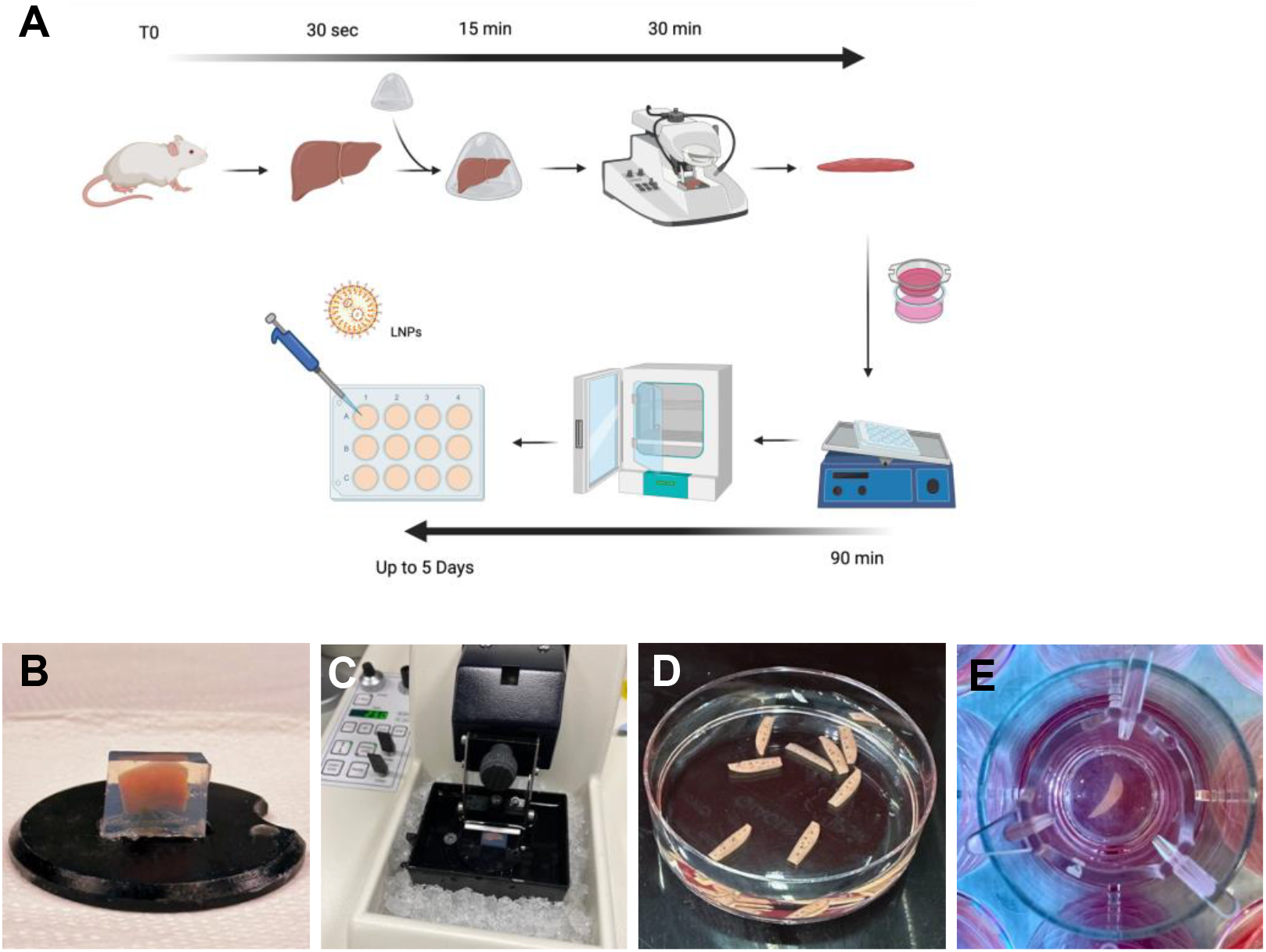
Optimised protocol for cutting and culture of PCLS. **(A)** Schematic summarising the protocol for generating PCLS. **(B)** Pre-cut liver lobe embedded in a low-melting agarose block. **(C)** Agarose block containing the mouse lobe ready for cutting onto vibratome filled with ice and ice-cold Krebs buffer. **(D)** PCLS following cutting and ready for culture. **(E)** A PCLS inside a transwell within a 12 well plate containing culture media.

### An optimised protocol of PCLS culture shows viability over 5 days

To determine the viability of PCLS, cell viability was assessed by 4,5-dimethylthiazol-2-yl)-5- (3-carboxymethoxyphenyl)-2-(4-sulfophenyl)-2H-tetrazolium (MTS) assay, which requires NAD(P)H-dependent dehydrogenases *i.e*. metabolically active cells, to reduce MTS. To optimise PCLS viability, we observed that a minimal volume of culture medium was essential to sustain viability after 24 hours of incubation. Higher volumes have been suggested to provide more nutrients but can dilute toxic bile acid products [44]. A volume of 0.7 mL in 24 well plate showed a significant reduction of viability by TMS assay (*p*=0.02) compared to 1.5 mL in 12 well plate and 2.6 mL in 6 well plate (Figure 2A). In agreement with others [45], we used 12 well plates as the best compromise for optimal survival within a smaller volume of culture medium.

**FIGURE 2.**
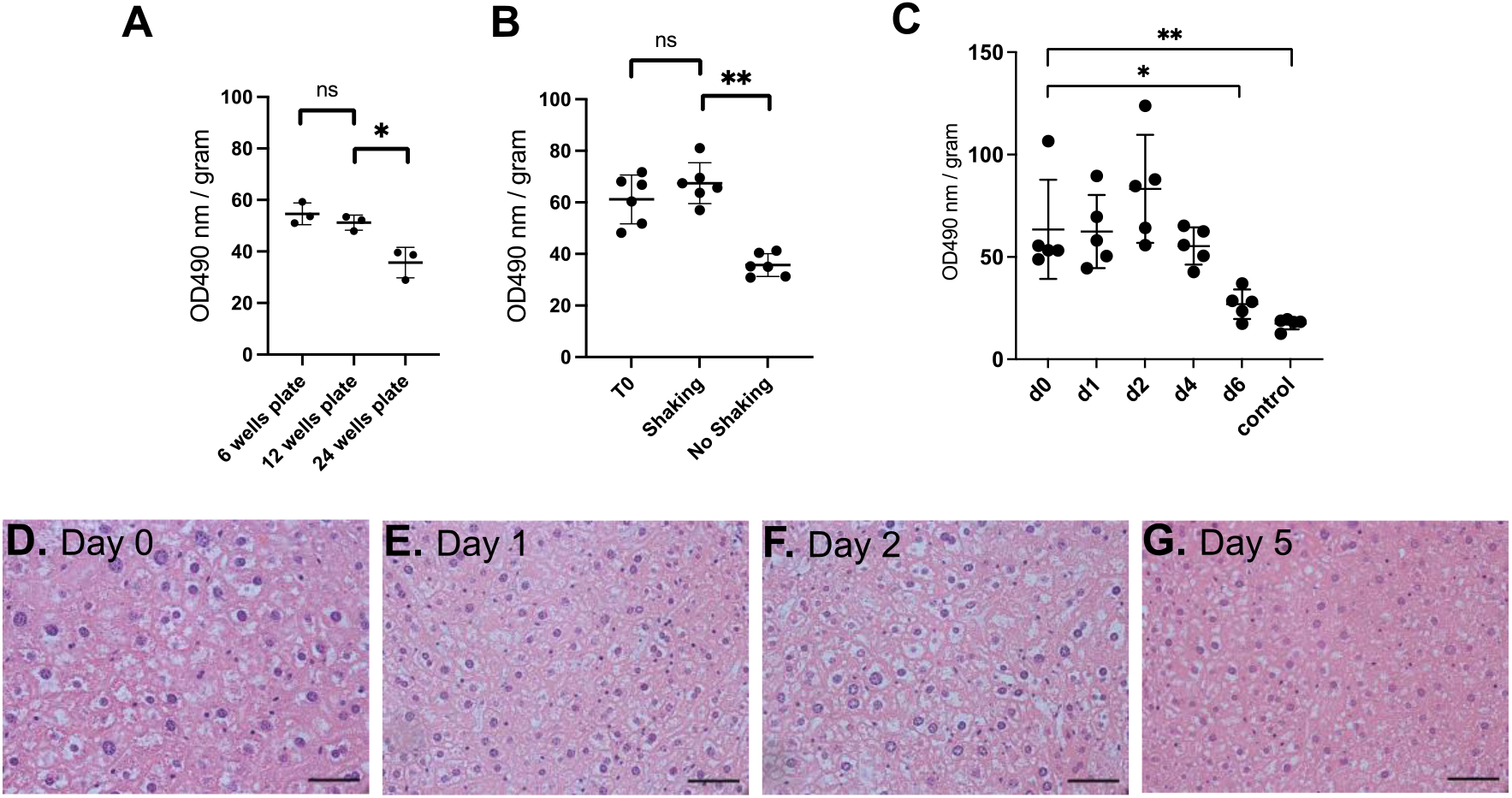
An optimised protocol of PCLS culture shows satisfactory viability for 5 days. **(A)** Effect of well size on cell viability (n=3). **(B)** Effect of shaking on cell viability (n=6 per condition). **(C)** MTS cell viability assay from liver sections from baseline until 6 days of incubation (n=5 per timepoint). OD: arbitrary unit of optical density, normalised to slice fresh weight. Graphs show mean ± SD. Unpaired 2-tailed Student’s t test, ns=not significant, *p<0.05, **p<0.01. **(D-G)** Representative images of histology of liver PCTS following H&E staining (n=3). Scale bar= 100 μM.

A 50% reduction of PCLS viability was observed at 24h post-incubation without continuous gentle shaking (Figure 3B). Shaking creates a critical air-liquid interface, optimised with the use of transwells allowing access to nutrients and oxygen to both faces of the section. Uptake of oxygen and nutrients is also increased by the constant flow created by the shaking movement but also passing through the transwell membrane [25, 46, 47].

**FIGURE 3.**
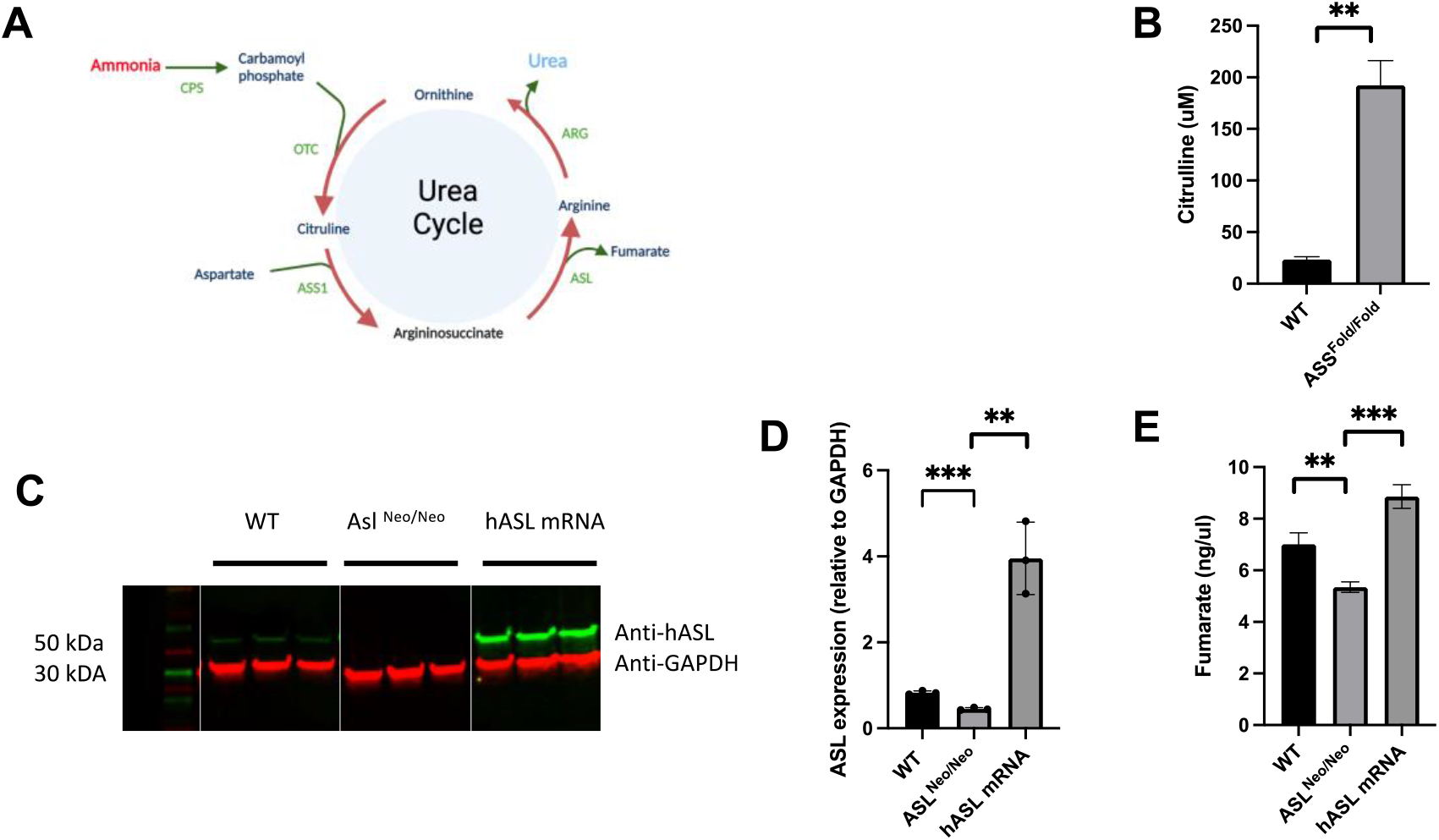
PCLS recapitulate the disease phenotype of ASS1 and ASL deficiencies and shows proof of concept of mRNA therapy *ex vivo*. **(A)** ASL and ASS1 enzymes enable ammonia detoxification in the liver-based urea cycle. **(B)** Citrulline levels in media after 48h of incubation in WT and ASS1-deficient PCLS. **(C)** ASL western blot at 48 hours. **(D)** Quantification of ASL immunoblot normalised to GAPDH. **(E)** Liver ASL activity from WT, untreated ASL^Neo/Neo^ and hASL mRNA. Graphs show mean ±SD. Unpaired 2-tailed Student’s t test, **p<0.01, ***p<0.005. **(B)** n=2 per group; **(C-E)** n=3 per group.

With these modifications, the MTS assay was assessed after an incubation from 1 hour up to 6 days. Cell viability remained constant for 5 days, from day 0 to 4 post-incubation before observing a significant decrease (*p*=0.05) at day 6 compare to day 0(Figure 2C). The PCLS morphology assessed by haematoxylin and eosin (H&E) staining showed no change of bile ducts and architecture up to 5 days post-incubation (Figures 2D-G). Compared to day 0 (Figure 2D), PCLS showed no histological differences at day 1 (Figure 2E) and 2 (Figure 2F) post incubation, with nuclear hyperchromasia, mild inflammatory infiltrate, vacuolisation in favour of a moderate cell death process at day 5 post-incubation (Figure 2G). Taken together, we showed that our optimised PCL S culture protocol enabled viability for 5 days.

### PCLS recapitulate the disease phenotype of ASS and ASL deficiencies and show proof of concept of mRNA replacement therapy *ex vivo*

Citrullinemia type 1 and argininosuccinic aciduria (ASA) are liver inherited metabolic diseases caused by defective ureagenesis and impaired clearance of neurotoxic ammonia. Citrullinemia type 1 and argininosuccinic aciduria (ASA) are caused by deficiency of the hepatic urea cycle enzymes argininosuccinate synthetase (ASS1) and argininosuccinate lyase (ASL), respectively (Figure 3A). Patients suffering from these urea cycle disorders develop hyperammonaemia, which causes acute coma and death if not aggressively treated in emergency with high dose of intravenous ammonia scavenger drugs and hemofiltration. Despite chronic standard of care based on ammonia scavengers and protein-restricted diet, patients present with high rates of mortality, neurological sequelae and poor quality of life [48]. The only cure is liver transplantation, which requires lifelong immunosuppression, has its own morbidity and is limited by organ shortage. Due to high unmet needs, novel therapies with small molecules [49] or gene therapies [50–53] are being developed. The *ex vivo* models rely on iPSC-derived hepatocytes with a relative inaccuracy in modelling the disease phenotype partially due to sub-optimal differentiation [3]. We therefore established a PCLS model that could provide the *ex vivo* characterisation of such metabolic defects by isolating and culturing PCLS from the hypomorphic *Ass^fold/fold^* mouse model of citrullinemia type 1, and the hypomorphic *Asl^Neo/Neo^* mouse recapitulating ASA. Similarly, the *Ass^fold/fold^* mouse model is characterised by impaired growth, abnormal fur, hyperammonaemia and high citrulline levels reproducing the clinical phenotype of citrullinemia type 1 [54]. The *Asl^Neo/Neo^* mice shows a similar phenotype with increased severity and early death at 3 weeks of age [55].

We generated PCLS from *Ass^fold/fold^* and wild-type (WT) littermate mice and harvested media at 48h to investigate citrulline levels, a key biomarker in ASS deficiency. We confirmed a significant 8-fold increase of citrulline levels in the media of *Ass^fold/fold^* PCLS compared to that of WT littermates (*p*=0.01) reflecting the phenotype observed in both the *Ass^fold/fold^* mice and patients (Figure 3B).

Next, we wanted to determine if PCLS can be used for assessing gene therapies. mRNA encapsulated in lipid nanoparticles is an attractive therapeutic strategy, which is rapidly being developed for rare liver inherited metabolic diseases [51, 56] and is in phase I/II clinical trials for 2 inherited metabolic diseases *i.e*. ornithine trancarbamylase deficiency (NCT04442347), the most common urea cycle disorder, propionic acidaemia (NCT04899310), methylmalonic acidaemia (NCT04159103) and glycogen storage disease 1A (NCT05095727) [57]. We generated PCLS from *Asl^Neo/Neo^* mice that we treated *ex vivo* with either *hASL* mRNA or PBS, compared to PCLS from WT littermates. All particles were added to the upper side of the slice to avoid a diluting effect into the culture medium as suggested by Griffin *et al* [58]. We assessed efficacy by assessing ASL expression and function by western blot and enzymatic activity at 48h post-transfection. An 8-fold increase in ASL protein levels was observed in *hASL* mRNA treated group compared to untreated PCLS, reaching supraphysiological ASL levels (Figures 3C, D). ASL degrades argininosuccinate into arginine and fumarate. ASL activity assessed by fumarate production was also restored in *hASL* mRNA versus PBS-treated PCLS to supraphysiological levels (Figure 3E). Taken together, these results demonstrate that PCLS culture yielded viable liver tissues replicating the biological phenotype observed in genetically modified mouse models. The PCLS model was amenable to successfully test therapeutic efficacy exemplified by non-viral gene therapy.

## Discussion

We demonstrate that PCLS can be an appealing *ex vivo* model to assess biological phenotype and therapeutic efficacy, whilst combining the advantage of respecting the complex liver architecture and reducing the use of animals.

Our method exemplifies a simple PCLS culture protocol with easy set-up, which requires minimal equipment. Our model can be replicated in a standard cell culture laboratory, which has access to an animal facility and a vibratome. Some previously published protocols required complex culture media and oxygen concentration higher than 80%, upregulating the metabolism and providing longer viability [9, 34, 36, 46]. Our experience shows this is not an absolute requirement. Moreover, the high oxygen concentration is likely to substantially modify the pathophysiology and the phenotype by generating toxic reactive oxygen species and subsequent antioxidant responses and signalling cascades [59]. Despite simple settings, our work emphasizes the importance of optimal vibratome settings, culture volumes and shaking. Liver slicing without a 3D supporting agarose gel results in damaged slices, uneven thickness and cell death. The use of hydrogel will have an improved benefit [60]. This optimisation allowed a viability maintained for 5 days, an acceptable compromise compared to the PCLS viability previously described between 48h to 10 days [38].

PCLS are essentially used to assess preclinical toxicity or pharmacokinetics [38, 61]. No attempt to replicate a biological phenotype of liver monogenic metabolic diseases had been reported previously. We expanded the application of this model by demonstrating that PCLS can replicate some key characteristics of Citrullinemia type 1 and ASA. This broadens the use of PCLS as a reliable *ex-vivo* model for preclinical studies assessing therapeutic screening for liver inherited metabolic diseases. Our approach could benefit other rare or common liver diseases, non-alcoholic fatty liver disease (NAFLD) [62], liver cancer [63] or even Fah^-/-^ Rag2^-/-^ Il2rg^-/-^ (FRG) mice with a chimeric humanised liver [64]. Liver PCLS have been used to study pathogenesis of hepatitis virus [65] or uptake of nanoparticles [66] but without gene therapy application.

The main limitations associated with PCLS are the short-lived culture system which last more than some days at best. The penetration of the therapeutic agent in the inner cell layers of the PCLS has been questioned [38]. Although this warrants further investigation, our experience shows that the efficacy endpoints tested in thin PCLS are satisfactory, thus enabling supraphysiological correction.

In conclusion, we show that PCLS can be developed with limited equipment and pioneer its use as a reliable *ex vivo* model for liver monogenic diseases, especially inherited metabolic diseases such as urea cycle defects. We show proof of concept that this model can be successfully developed for assessing therapeutic efficacy of gene therapy. We believe this model provides *ex vivo* high-throughput possibilities whilst preserving the tissue complexity and should therefore become a more recognised tool for preclinical studies in rare and common liver diseases.

## Methods and Protocols

### mRNA formulation

*hASL* and Luciferase (*Luc*) encoding mRNA encapsulated in Lipid Nanoparticles (LNPs) were provided by Moderna Therapeutics using their proprietary technology. Codon optimized mRNA encoding *hASL* was synthesized in vitro by T7 RNA polymerase-mediated transcription. The mRNA initiated with a cap, followed by a 5’ untranslated region (UTR), an open reading frame (ORF) encoding *hASL*, a 3’ UTR and a polyadenylated tail. Uridine was globally replaced with N1-methylpseudouridine, previously described [67]. For in vivo intravenous delivery, LNP formulations were generated. Briefly, mRNA was mixed with lipids at a molar ratio of 3:1 (mRNA: lipid), previously described [68]. mRNA-loaded nanoparticles were exchanged into final storage buffer and had particle sizes of 80 - 100 nm, >80% encapsulation of the mRNA by RiboGreen assay, and <10 EU/mL endotoxin levels.

### Animals

The following homozygous transgenic mouse strains purchased from Jackson Laboratory (Bar Harbor, ME) were used in this study: *Asl^Neo/Neo^ (B6.129S7-Asltm1Brle/J*) and Ass1^fold^ (*B6EiP-Ass1^fold^/GrsrJ*). Livers from littermates were harvested between the ages of day 12 and 19. Animals were maintained on standard rodent chow (Harlan 2018, Teklab Diets, Madison, WI) with free access to water in a 12-hour light / 12 hours dark environment. WT littermates were used as controls and housed in the same cages. Genotyping was performed using DNA extracted from tail clips as previously described [69]. Animal procedures were performed under the UK Home Office licences 70/8030 and PP9223137, compliant with ARRIVE guidelines.

### Collection of liver and preparation

The liver was excised as from the mouse and in conditions as sterile as possible to minimise the risk of contamination and store in ice cold Krebs Buffer. All further steps were performed on ice at 4°C. Each lobe was isolated from the whole liver and all edges further trimmed to obtain a smaller more manageable lobe with straight edges. This helped removing some of the fibrous Glisson’s capsule to further facilitate sectioning. This was performed while keeping the liver surfaces wet into ice cold Krebs buffer (Fig 1A). 1L of KREBS (Merck, cat. no K3753) buffer was prepared by dissolving one vial of Krebs powder into 1L of ultrapure water and kept at 4°C before and during use. Each lobe section was embedded into 4% low melting agarose (ThermoFisher, Cat no 16520050). The agarose was let cool down and cut it further into more manageable cubes ready for sectioning. The agarose cubes was kept on ice cold Krebs Buffer to preserve the tissues in cold conditions until sectioning.

### Liver slices preparation

Slicing was performed using a vibratome (Leica, VT1000 S). The blades (Agar Scientific, cat no T569T) were placed onto the vibratome at an angle of 10 degree downwards and below horizontal (Fig 1C). The vibratome was set for cutting at a thickness of 250 μm and speed was set at 5 and frequency at 7. The cutting tray was cooled into the freezer before use and preserved cold with ice around it (Fig 1C). The vibratome platform and blades were disinfected with 70% ethanol followed by a rinse with a saline solution. The agarose blocks were glued, using standard cyanoacrylate glue, directly onto the platform and the tray filled with ice cold Krebs buffer to completely cover the agarose block ready for sectioning.

The lobe being positioned and cut in a transversal position prevented further damage to the tissue. A spatula was used to collect the liver slices instead of forceps or brushes to avoid damaging the slices.

### Incubation of liver slices

William’s Medium E with GlutaMAX™ (WME) slice incubation medium was prepared by adding 2mM L-glutamine supplement (Gibco, cat no 32551-020), 10% of dialysed FBS (ThermoFisher, cat no 26400044), 100 U/mL penicillin and 100 μg/mL streptomycin (Gibco, Cat no 15750045), 10μg/mL Gentamycin (Gibco, Cat no 15750045), 25mM D-Glucose solution (Gibco, Cat no 15384895), 15mM HEPES solution (Gibco, Cat no 15630), stored at 4°C. The slices were transferred, using a spatula to avoid damage, into a 12 well plates containing 1.5 ml of prewarmed complete WME media. The quality sections was judged based on smooth edges and even thickness. Slices were also culture using porous 8μm inserts (Strastedt, Cat no 83.3932.800). Plates with slices were transferred into a humidified incubator set to 37°C, 5% carbon dioxide and 20% normoxic oxygen level while continuously shaking using an orbital shaker with a speed set at 130rpm to allow optimal mixing of nutrients and increased oxygenation. Thereafter the media was changed every 48h. The plate needed to be protective from the shaker’s platform to prevent overheating of the culture media.

### MTS Assay

All MTS Assay were performed using livers from wild type *Bl6* mice. The slices were transferred into a 48 well plate containing 400 μl of prewarmed complete WME media and 80ul of MTS tetrazolium reagent (Abcam, Cat no ab197010) was added to generate a formazan dye. Following incubation for 1 h at 37°C, 5% CO_2_ onto a shaker, 200 μl of media was transferred into a 96 well plate and absorbance was measured at 490 nm using a multiwell plate reader (Promega). Slices left on the bench in water and at room temperature for 24h were used as negative controls.

### Histology

Slices were fixed in 10% Formalin, neutral buffered (Merck, Cat No HT501128) for 48h and transferred into 70% Ethanol before being embedded in paraffin and cut into 4μm sections and stained with haematoxylin and eosin using standard protocol by the Pathology lab, UCL Institute of Neurology. The slides were imaged under Zeiss Axioplan Histology scope at UCL Great Ormond Street Institute of Child Health Imaging Facility.

### Protein Assay

Slices samples were immediately transferred following collection onto dry ice and kept at −80°C until analysis. Tissues were homogenised into 200 μl of ice-cold 1x RIPA buffer (Cell signalling) using a bead homogenesiser (Qiagen). The samples were centrifuged at 10000g for 20 min at 4°C and protein-containing supernatant transferred into a new tube. The protein concentration was measured using a BCA assay kit (ThermoFisher) per manufacturer’s instructions.

### Western Blot

50 μl samples containing 50 μg total protein per sample was diluted with 4x Laemmli sample buffer (containing 10% 2-β-mercaptoethanol) at a final volume of 50ul, vortexed and heated to 95°C for 10 min. SDS-PAGE was used to separate the proteins at 100V for 1h followed by wet transfer of proteins into an immobilin PVDF membrane at 400mA for 1h. Blocking was performed with 5% non-fat milk in PBS-T (1×PBS with 0.1% tween-20) for one hour at room temperature and subsequently incubated overnight at 4°C with primary antibodies (Anti-ASL, Abcam ab97370, 1:1,000; Anti-GAPDH mouse, Abcam ab8245, 1:10,000). The membranes were then washed three times in PBS-T and incubated with secondary antibodies (Anti-ASL, Abcam ab97370, 1:1,000; Anti-GAPDH mouse, Abcam ab8245, 1:10,000; Anti-nitrotyrosine, Merck 05-233, 1:100; Anti-GAPDH rabbit, Abcam ab9485, 1:1,000), 3x 5-min washes with PBS-T, 1h incubation with fluorescent secondary antibodies (IRDye^®^ 800CW Goat anti-Rabbit IgG 1:1,000, 926-32210 and IRDye^®^ 680RD Donkey anti-Mouse IgG, 923-68072, Licor) and 3x 5 min washes with PBS-T. Image acquisition and analysis was performed using Licor Odyssey and image analysed using Licor ImageStudio Lite software. ASL protein signal per sample was normalised against GAPDH which was used as a loading control.

### ASL enzyme activity

For liver ASL activity, 20-30mg of liver was homogenised in 400μL of cold homogenising buffer (50mM phosphate buffer pH 7.5 and 1x Roche EDTA-free protease inhibitor (Roche, Switzerland)) using Precellys homogeniser tube (VWR, UK) and Precellys 24 tissue homogeniser (Bertin Instruments, France), centrifuged at 10000g for 20 min at 4°C and protein levels measured from the supernatant using BCA kit (Thermo Fisher Scientific, UK). 60μg of protein lysate was incubated with 3.6mM ASA in final volume of 50μl, incubated at 37°C for 1 h followed by reaction termination at 80°C for 20 min. The mixture was centrifuged at 10000g for 5min and 5μL of the supernatant was used to measure fumarate levels per instruction from the commercial fumarate kit (Abcam, Cambridge, UK).

### Nanoparticle transfection

Lipid Nanoparticles containing mRNA (LNP-hASL mRNA) all expressing human ASL were added to the upper side of the slice. 2 μg of LNP-hASL in a 10μL volume were added to each slice. Slices and media were harvested at 48h. PCLS were produced and cultured in parallel from livers originating from aged matched litter mate wildtype controls.

### Citrulline analysis

40μl media at various timepoints was mixed with 30μl of 0.1M HCl and 10ul of IS mixture (final concentration of 2nmoL). The sample was then topped up with 280μL of Solvent A, vortexed and incubated at room temperature for 5 minutes following by centrifugation at 16,000 rpm for 5 minutes at room temperature. The supernatant was transferred on to glass vials for separation and detection of amino acids, performed by hydrophilic interaction Liquid chromatography (HILIC) coupled with tandem mass spectrometry, a method adapted from [70]. Amino acid chromatography separation was performed in Acquity UltraPure Liquid Chromatography (UPLC)-system (Waters, Manchester, United Kingdom) using Acquity UPLC BEH Amide column (2.1×100mm, 1.7μm particle size) including a Van Guard™ UPLC BEH Amide pre-column (2.1×5mm, 1.7μm particle size) (Waters Limited, UK). The mobile phases were (A) 10mM ammonium formiate in 85% acetonitrile (A0627/17, Fisher Scientific) and 0.15% formic acid (A117-50, FisherScientific), (B) 15mM ammonium formiate (Sigma-Aldrich) in MilliQ-water containing 0.15% formic acid, pH 3.0. Initial conditions were 100% solvent A, flow rate of 0.4ml/min. At 6.1 minutes, solvent A decreased to 94.1% and solvent B increased to 5.9%, then at 10.1 minutes, solvent A was further decreased to 82.4% and solvent B to 17.6%. At 13.1 minutes, solvent A was further decreased to 70.6% and solvent B increased to 29.4% with flow rate of 0.6ml/min. At 17.1 minutes, solvent A was increased back to 100% with flow rate of 0.4ml/min. At 18.6 minutes, combination of 50% solvent A and 50% solvent B was applied at 0.4ml/min as a wash step to remove any carryovers. The column was then re-equilibrated for an additional 10 minutes with initial starting conditions. Detection was performed using a tandem mass spectrometer Xevo TQ-S (Waters, UK) using multiple reaction monitoring in positive ion mode. The dwell time was set automatically with MRM-transition for citrulline 176.15>159.05 and L-citrulline-d7 of 183.15>166.05. L-Citrulline-d7 (CDN Isotopes, Cambridge) was used as internal standard control for ASA and their anhydrides. Cone voltage and collision energy of 25V was used for both ASA and their anhydrides while cone voltage and collision energy of 15V and 10V respectively was used for L-citrulline-d7. The source temperature was 150°C, desolvation temperature 550°C, capillary voltage 1.00 kV, cone gas flow 150 L/hour and desolvation gas flow 500 L/h. Data was acquired using Masslynx v4.2 software (Waters, UK). TargetLynx™ (Waters, UK) application manager was used for subsequent batch data processing and reporting of results. Calibration curve of unlabelled ASA were used for quantification of 15N-ASA.

### Statistical Analyses

GraphPad Prism 9.0 software (San Diego, CA, USA) was used for performing data analysis and generating graphs.

## Supplementary Material

None

## Acknowledgements

The authors thank Rebecca Towns, Mirabela Bandol, Samantha Richards, Katherine Howett, Louise Fisher and the staff from UCL Biological Services for their help with breeding and maintenance of the animal colonies.

## Author Contributions

DP and JB designed the work. DP conducted the experimental work. GS, LT, SG, CD provided technical assistance in experimental work. SG conducted mass spectrometry analysis. DP wrote the manuscript. All authors revised substantially and approved the final version of the manuscript. This manuscript has not been accepted or published elsewhere.

## Institutional review board statement

Animal procedures were performed under the UK Home Office licences 70/8030 and PP9223137.

## Data Availability Statement

Data are available upon request to the corresponding author.

## References

1. Dong, L., et al., The role of the microenvironment on the fate of adult stem cells. Sci China Life Sci, 2015. 58(7): p. 639–48.

2. Ferguson, L.P., E. Diaz, and T. Reya, The Role of the Microenvironment and Immune System in Regulating Stem Cell Fate in Cancer. Trends Cancer, 2021. 7(7): p. 624–634.

3. Duff, C. and J. Baruteau, Modelling urea cycle disorders using iPSCs. NPJ Regen Med, 2022. 7(1): p. 56.

4. Lorvellec, M., et al., An In Vitro Whole-Organ Liver Engineering for Testing of Genetic Therapies. iScience, 2020. 23(12): p. 101808.

5. ; Available from: https://www.nc3rs.org.uk.

6. Mondonedo, J.R., et al., A High-Throughput System for Cyclic Stretching of Precision-Cut Lung Slices During Acute Cigarette Smoke Extract Exposure. Front Physiol, 2020. 11: p. 566.

7. Viana, F., C.M. O’Kane, and G.N. Schroeder, Precision-cut lung slices: A powerful ex vivo model to investigate respiratory infectious diseases. Mol Microbiol, 2022. 117(3): p. 578–588.

8. Iulianella, A., Cutting Thick Sections Using a Vibratome. Cold Spring Harb Protoc, 2017. 2017(6): p. pdb prot094011.

9. Pearen, M.A., et al., Murine Precision-Cut Liver Slices as an Ex Vivo Model of Liver Biology. J Vis Exp. 2020 (157).

10. De Kanter, R., et al., Prediction of whole-body metabolic clearance of drugs through the combined use of slices from rat liver, lung, kidney, small intestine and colon. Xenobiotica, 2004. 34(3): p. 229–41.

11. van de Kerkhof, E.G., et al., Induction of metabolism and transport in human intestine: validation of precision-cut slices as a tool to study induction of drug metabolism in human intestine in vitro. Drug Metab Dispos, 2008. 36(3): p. 604–13.

12. Nogueira, G.O., et al., Modeling the Human Brain With ex vivo Slices and in vitro Organoids for Translational Neuroscience. Front Neurosci, 2022. 16: p. 838594.

13. Ucar, B., N. Stefanova, and C. Humpel, Spreading of Aggregated alpha-Synuclein in Sagittal Organotypic Mouse Brain Slices. Biomolecules, 2022. 12(2).

14. Liu, G., et al., Precision cut lung slices: an ex vivo model for assessing the impact of immunomodulatory therapeutics on lung immune responses. Arch Toxicol, 2021. 95(8): p. 2871–2877.

15. Klouda, T., et al., Precision Cut Lung Slices as an Efficient Tool for Ex vivo Pulmonary Vessel Structure and Contractility Studies. J Vis Exp. 2021 (171).

16. De Kanter, R., et al., Drug-metabolizing activity of human and rat liver, lung, kidney and intestine slices. Xenobiotica, 2002. 32(5): p. 349–62.

17. Stribos, E.G.D., et al., Murine Precision-Cut Kidney Slices as an ex vivo Model to Evaluate the Role of Transforming Growth Factor-beta1 Signaling in the Onset of Renal Fibrosis. Front Physiol, 2017. 8: p. 1026.

18. James, K., G. Skibinski, and P. Hoffman, A comparison of the performance in vitro of precision cut tissue slices and suspensions of human spleen with special reference to immunoglobulin and cytokine production. Hum Antibodies Hybridomas, 1996. 7(4): p. 138–50.

19. Tatibana, M., K. Kita, and T. Asai, Stimulation by 6-azauridine of carbamoyl phosphate synthesis for pyrimidine biosynthesis in mouse spleen slices. Eur J Biochem, 1982. 128(2-3): p. 625–9.

20. Camelliti, P., et al., Adult human heart slices are a multicellular system suitable for electrophysiological and pharmacological studies. J Mol Cell Cardiol, 2011. 51(3): p. 390–8.

21. Liu, Z., et al., Comparative analysis of adeno-associated virus serotypes for gene transfer in organotypic heart slices. J Transl Med, 2020. 18(1): p. 437.

22. Zimmermann, M., et al., Human precision-cut liver tumor slices as a tumor patient-individual predictive test system for oncolytic measles vaccine viruses. Int J Oncol, 2009. 34(5): p. 1247–56.

23. Philouze, P., et al., CD44, gamma-H2AX, and p-ATM Expressions in Short-Term Ex Vivo Culture of Tumour Slices Predict the Treatment Response in Patients with Oral Squamous Cell Carcinoma. Int J Mol Sci, 2022. 23(2).

24. Tigges, J., et al., Optimization of long-term cold storage of rat precision-cut lung slices with a tissue preservation solution. Am J Physiol Lung Cell Mol Physiol, 2021. 321(6): p. L1023-L1035.

25. Olinga, P., et al., Comparison of five incubation systems for rat liver slices using functional and viability parameters. J Pharmacol Toxicol Methods, 1997. 38(2): p. 59–69.

26. Ou, Q., et al., Physiological Biomimetic Culture System for Pig and Human Heart Slices. Circ Res, 2019. 125(6): p. 628–642.

27. Martin, C., Human Lung Slices: New Uses for an Old Model. Am J Respir Cell Mol Biol, 2021. 65(5): p. 471–472.

28. Sewald, K. and O. Danov, Infection of Human Precision-Cut Lung Slices with the Influenza Virus. Methods Mol Biol, 2022. 2506: p. 119–134.

29. Lerche-Langrand, C. and H.J. Toutain, Precision-cut liver slices: characteristics and use for in vitro pharmaco-toxicology. Toxicology, 2000. 153(1-3): p. 221–53.

30. https://www.nc3rs.org.uk. Accessed 08/02/2023.

31. https://www.gov.uk/government/publications/liver-disease-applying-all-our-health/liver-disease-applying-all-our-health#fn:3. Accessed 08/02/2023.

32. Moon, A.M., A.G. Singal, and E.B. Tapper, Contemporary Epidemiology of Chronic Liver Disease and Cirrhosis. Clin Gastroenterol Hepatol, 2020. 18(12): p. 2650–2666.

33. Baruteau, J., et al., Gene therapy for monogenic liver diseases: clinical successes, current challenges and future prospects. J Inherit Metab Dis, 2017. 40(4): p. 497–517.

34. de Graaf, I.A., et al., Preparation and incubation of precision-cut liver and intestinal slices for application in drug metabolism and toxicity studies. Nat Protoc, 2010. 5(9): p. 1540–51.

35. Koch, A., et al., Murine precision-cut liver slices (PCLS): a new tool for studying tumor microenvironments and cell signaling ex vivo. Cell Commun Signal, 2014. 12: p. 73.

36. Szalowska, E., et al., Effect of oxygen concentration and selected protocol factors on viability and gene expression of mouse liver slices. Toxicol In Vitro, 2013. 27(5): p. 1513–24.

37. Brugger, M., et al., High precision-cut liver slice model to study cell-autonomous antiviral defense of hepatocytes within their microenvironment. JHEP Rep, 2022. 4(5): p. 100465.

38. Dewyse, L., H. Reynaert, and L.A. van Grunsven, Best Practices and Progress in Precision-Cut Liver Slice Cultures. Int J Mol Sci, 2021. 22(13).

39. Smith, J.T., et al., Effect of slice thickness on liver lesion detection and characterisation by multidetector CT. J Med Imaging Radiat Oncol, 2010. 54(3): p. 188–93.

40. Starokozhko, V., et al., Maintenance of drug metabolism and transport functions in human precision-cut liver slices during prolonged incubation for 5 days. Arch Toxicol, 2017. 91(5): p. 2079–2092.

41. Cascarano, J., W.L. Chick, and B.W. Zweifach, Glycogen synthesis in rat liver slices: role of glucose concentration, mesothelium and insulin. Proc Soc Exp Biol Med, 1966. 121(2): p. 619–22.

42. Salans, L.B. and G.M. Reaven, Effect of insulin pretreatment on glucose and lipid metabolism of liver slices from normal rats. Proc Soc Exp Biol Med, 1966. 122(4): p. 1208–13.

43. Roden, M. and E. Bernroider, Hepatic glucose metabolism in humans--its role in health and disease. Best Pract Res Clin Endocrinol Metab, 2003. 17(3): p. 365–83.

44. Graaf, I.A., G.M. Groothuis, and P. Olinga, Precision-cut tissue slices as a tool to predict metabolism of novel drugs. Expert Opin Drug Metab Toxicol, 2007. 3(6): p. 879–98.

45. van de Kerkhof, E.G., et al., Characterization of rat small intestinal and colon precision-cut slices as an in vitro system for drug metabolism and induction studies. Drug Metab Dispos, 2005. 33(11): p. 1613–20.

46. Leeman, W.R., I.A. van de Gevel, and A.A. Rutten, Cytotoxicity of retinoic acid, menadione and aflatoxin B(1) in rat liver slices using Netwell inserts as a new culture system. Toxicol In Vitro, 1995. 9(3): p. 291–8.

47. Schumacher, K., et al., Perfusion culture improves the maintenance of cultured liver tissue slices. Tissue Eng, 2007. 13(1): p. 197–205.

48. Baruteau, J., et al., Expanding the phenotype in argininosuccinic aciduria: need for new therapies. J Inherit Metab Dis, 2017. 40(3): p. 357–368.

49. Soria, L.R., et al., Beclin-1-mediated activation of autophagy improves proximal and distal urea cycle disorders. EMBO Mol Med, 2021. 13(2): p. e13158.

50. Touramanidou, L., Gurung S, Cozmescu AC, Waddington SN, Counsell JR, Gissen P, Baruteau J., In vivo lentiviral gene therapy for argininosuccinic aciduria. European Society of Gene and Cell Therapy (ESGCT); Virtual Congress: Human Gene Therapy, 2021. 32: p. A1-152.

51. Gurung, S., Timmermand, O.V., Perocheau P, Gil-Martinez AL, Minnion M, Touramanidou L, Fang S, Messina M, Khalil Y, Barber AR, Edwards RS, Finn PF, Cavedon A, Siddiqui S, Rice L, Martini PGV, Mills PB, Waddington SN, Gissen P, Eaton S, Ryten M, Feelisch M, Frassetto A, Witney TH*, Baruteau J*. mRNA therapy restores ureagenesis and corrects glutathione metabolism in argininosuccinic aciduria. BioRxiv, 2022. https://biorxiv.org/cgi/content/short/2022.10.19.512931v1.

52. Baruteau, J.P., D. P.; Hanley, J.; Lorvellec, M.; Rocha-ferreira, E.; Karda, R.; Ng, J.; Suff, N.; Antinao Diaz, J.; Rahim, A. A.; Hughes, M. P.; Banushi, B.; Prunty, H.; Hristova, L.; Ridout, D. A.; Virasami, A.; Heales, S.; Howes, S. J.; Buckley, S. M.; Mills, P. B.; Gissen, P.; Waddington, S. N., Argininosuccinic aciduria fosters neuronal nitrosative stress reversed by Asl gene transfer. Nature Communications, 2018. In Press.

53. Kok, C.Y., et al., Adeno-associated virus-mediated rescue of neonatal lethality in argininosuccinate synthetase-deficient mice. Mol Ther, 2013. 21(10): p. 1823–31.

54. Perez, C.J., et al., Two hypomorphic alleles of mouse Ass1 as a new animal model of citrullinemia type I and other hyperammonemic syndromes. Am J Pathol, 2010. 177(4): p. 1958–68.

55. Erez, A., et al., Requirement of argininosuccinate lyase for systemic nitric oxide production. Nat Med, 2011. 17(12): p. 1619–26.

56. Martini, P.G.V. and L.T. Guey, A New Era for Rare Genetic Diseases: Messenger RNA Therapy. Hum Gene Ther, 2019. 30(10): p. 1180–1189.

57. Clinicaltrials.gov. accessed 8th December 2022:[

58. Griffin, J.M., et al., Astrocyte-selective AAV gene therapy through the endogenous GFAP promoter results in robust transduction in the rat spinal cord following injury. Gene Ther, 2019. 26(5): p. 198–210.

59. t Hart, N.A., et al., Oxygenation during hypothermic rat liver preservation: an in vitro slice study to demonstrate beneficial or toxic oxygenation effects. Liver Transpl, 2005. 11(11): p. 1403–11.

60. Bailey, K.E., et al., Embedding of Precision-Cut Lung Slices in Engineered Hydrogel Biomaterials Supports Extended Ex Vivo Culture. Am J Respir Cell Mol Biol, 2020. 62(1): p. 14–22.

61. Smith, P.F., et al., Dynamic organ culture of precision liver slices for in vitro toxicology. Life Sci, 1985. 36(14): p. 1367–75.

62. Nagarajan, P., et al., Genetically modified mouse models for the study of nonalcoholic fatty liver disease. World J Gastroenterol, 2012. 18(11): p. 1141–53.

63. Pascale, R.M., et al., Experimental Models to Define the Genetic Predisposition to Liver Cancer. Cancers (Basel), 2019. 11(10).

64. Cabanes-Creus, M., et al., Restoring the natural tropism of AAV2 vectors for human liver. Sci Transl Med, 2020. 12(560).

65. Kartasheva-Ebertz, D., et al., Adult human liver slice cultures: Modelling of liver fibrosis and evaluation of new anti-fibrotic drugs. World J Hepatol, 2021. 13(2): p. 187–217.

66. Dragoni, S., et al., Gold nanoparticles uptake and cytotoxicity assessed on rat liver precision-cut slices. Toxicol Sci, 2012. 128(1): p. 186–97.

67. Nelson, J., et al., Impact of mRNA chemistry and manufacturing process on innate immune activation. Sci Adv, 2020. 6(26): p. eaaz6893.

68. An, D., et al., Systemic Messenger RNA Therapy as a Treatment for Methylmalonic Acidemia. Cell Rep, 2017. 21(12): p. 3548–3558.

69. Baruteau, J., et al., Argininosuccinic aciduria fosters neuronal nitrosative stress reversed by Asl gene transfer. Nat Commun, 2018. 9(1): p. 3505.

70. Prinsen, H., et al., Rapid quantification of underivatized amino acids in plasma by hydrophilic interaction liquid chromatography (HILIC) coupled with tandem mass-spectrometry. J Inherit Metab Dis, 2016. 39(5): p. 651–660.

